# Generative modeling of histology tissue reduces human annotation effort for segmentation model development

**DOI:** 10.1101/2021.10.15.464564

**Authors:** Brendon Lutnick, Pinaki Sarder

**Affiliations:** Department of Pathology & Anatomical Sciences, SUNY Buffalo, USA; Department of Biomedical Engineering, SUNY Buffalo, USA

## Abstract

Segmentation of histology tissue whole side images is an important step for tissue analysis. Given enough annotated training data modern neural networks are capable accurate reproducible segmentation, however, the annotation of training datasets is time consuming. Techniques such as human in the loop annotation attempt to reduce this annotation burden, but still require a large amount of initial annotation. Semi-supervised learning, a technique which leverages both labeled and unlabeled data to learn features has shown promise for easing the burden of annotation. Towards this goal, we employ a recently published semi-supervised method: *datasetGAN* for the segmentation of glomeruli from renal biopsy images. We compare the performance of models trained using *datasetGAN* and traditional annotation and show that *datasetGAN* significantly reduces the amount of annotation required to develop a highly performing segmentation model. We also explore the usefulness of using *datasetGAN* for transfer learning and find that this greatly enhances the performance when a limited number of whole slide images are used for training.

## INTRODUCTION

The segmentation of histological tissue structures from whole slide images (WSIs) is often an important first step for further downstream analysis and is therefore well explored in the literature [1-6]. The use of convolutional neural networks (CNNs) is currently considered the state of the art, however the generation of annotated training sets for segmentation of WSIs is time consuming and requires domain level expertise [7]. Semi-supervised learning has been successfully used to reduce the amount of annotation needed to generate performant models by utilizing information from unlabeled training images. While the adoption of semi-supervised learning to medical image datasets has lagged behind the field of computer science, this approach is ideal for histology data, where unlabeled data is plentiful.

Generally semi-supervised learning is accomplished by combining an unsupervised architecture such as a autoencoder [8] with a supervised cost function that is enforced only when labeled data is present [9]. The unsupervised architecture attempts to learn patterns in the data distribution, and is pushed to learn features important for classification by the influence of the supervised loss on parameter optimization [10]. Simply put, semi supervised learning attempts to leverage information learned from an unsupervised low dimensional representation of input images to inform a downstream classification task. However, recently the use of autoencoders has been surpassed by generative adversarial networks (GANs) due to their ability to reconstruct realistic looking images from low dimensional spaces [11]. GANs have been used to generate fake histology images but the usefulness of such applications is limited [12]. Unlike autoencoders, GANs focus on producing images, and have no way to encode the images into features, which is typically the direction of interest for classification tasks.

Recently, several approaches for utilizing GAN architectures for semi supervised segmentation training have been proposed [13, 14]. One promising method is Nvidia’s recently published *datasetGAN* [14], which proposes training a simple decoder network to predict semantic labels for synthetic images produced by the generator. Using this method the state of the art GAN architecture *styleGAN* [15] is first trained without labels, to produce realistic synthetic images. A decoder network is then be trained to produce an accurate pixelwise labeling of the synthetic images given a handful of labeled images. This allows the GAN to function as a labeled data factory, capable of producing large and heterogenous labeled datasets which can be used to train a supervised architecture such as *deepLab V3+* [16]. *DatasetGAN*, has shown compelling results for segmentation of natural images and seemed ideal for application to WSIs.

To test the feasibility of using *datasetGAN* to accelerate the development of models for segmentation of WSIs, we use it to train a model for glomeruli segmentation from renal tissue biopsies. We test the effectiveness of this method against traditional annotation and training using *Histo-Cloud* [4], our recently developed pipeline for cloud based WSI segmentation. Models created with *datasetGAN* require very minimal annotation effort, making them an ideal way to initialize network parameters for transfer learning. We therefore also test the ability of the model trained with *datasetGAN* annotations to improve further supervised training.

## RESULTS

To test the effectiveness of *datasetGAN* as a labeled data factory we compared the tradeoff between human annotation effort and glomeruli segmentation performance obtained. We trained and *deepLab V3+* network [16] using four different training sets to obtain four segmentation models. The first model was trained with a synthetic dataset of 50,000 image patches generated using *datasetGAN*, examples of synthetic data are shown in *Figure 1*. While the unsupervised training of the GAN was done using 2455 WSIs, only 24 synthetic glomeruli were hand annotated, examples of the annotated synthetic glomeruli are shown in *Figure 1*. When tested on a holdout dataset containing 23 WSIs the *datasetGAN* model produced a Mathews correlation coefficient (*MCC*) *= 0*.*68*. To test how this compares to a training using a hand annotated dataset, a *control* model was trained using four hand annotated WSIs which contained 76 labeled glomeruli. The control model performed worse than the *datasetGAN* model, resulting in an *MCC = 0*.*42* despite receiving more than 3X the annotation effort. To see if this performance would improve with the addition of extra training data, 5 more WSIs were added to the training set for a total of 161 annotated glomeruli, we all this the *control+* model. The addition of more training data did improve the performance of the *control+* model (*MCC = 0*.*61*), but still fell below the model trained using the *datasetGAN* dataset. Finally, to test the effectiveness transfer learning using *datasetGAN* we retrained the *datasetGAN* model using the *control* dataset. This significantly improved the performance of the *datasetGAN* model resulting in *MCC = 0*.*78*. The performance of the trained models is detailed in *Table 1*, and examples of the segmentation performance using the *datasetGAN* model is highlighted in *Figure 2*.

**Table 1.** Holdout segmentation performance using different training sets. The segmentation performance of the DeepLab v3+ network trained using various methods. The datasetGAN training set contained 24 annotated glomeruli, but was exposed to 2455 WSIs which were used to train styleGAN. This network performed better than using the control dataset (4 WSIs containing 76 annotated gloms), or the control+ dataset which added an additional 5 WSIs to the control set for a total of 161 annotated glomeruli. The most performant network was trained by retraining the datasetGAN on the control dataset using transfer learning.

| Training Set | Annotated Gloms | Training Slides | MCC |
| --- | --- | --- | --- |
| *DatasetGAN* | 24 | 2455 | 0.68 |
| Control | 76 | 4 | 0.42 |
| Control+ | 161 | 9 | 0.61 |
| Transfer learning<br>*DatasetGAN + Control* | 100 | 2455+4 | 0.78 |

**Figure 1.**
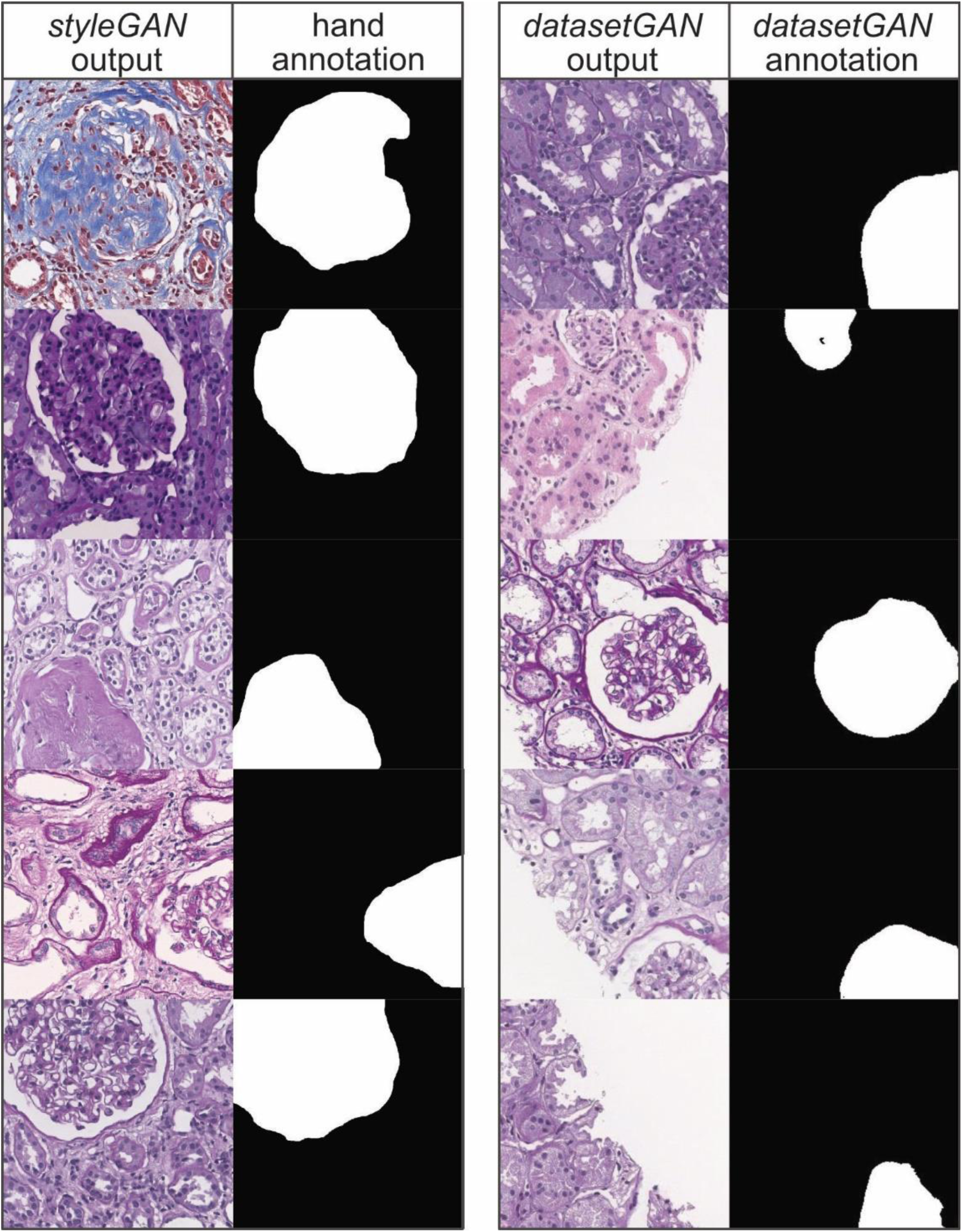
Examples of datasetGAN inputs and outputs. The panels on the left are examples of the hand annotations used to train datasetGAN. 24 selected styleGAN outputs were annotated for glomeruli by hand. The panels on the right are examples of labeled images produced by datasetGAN. The datasetGAN outputs are generated by the pre-trained styleGAN branch, and semantic annotations are automatically produced by the decoder branch of datasetGAN. Due to memory constraints when training datasetGAN, the outputs are produced at 1/4^*th*^ *the size of the originally trained styleGAN (256×256 pixels instead of 1024×1024)*.

**Figure 2.**
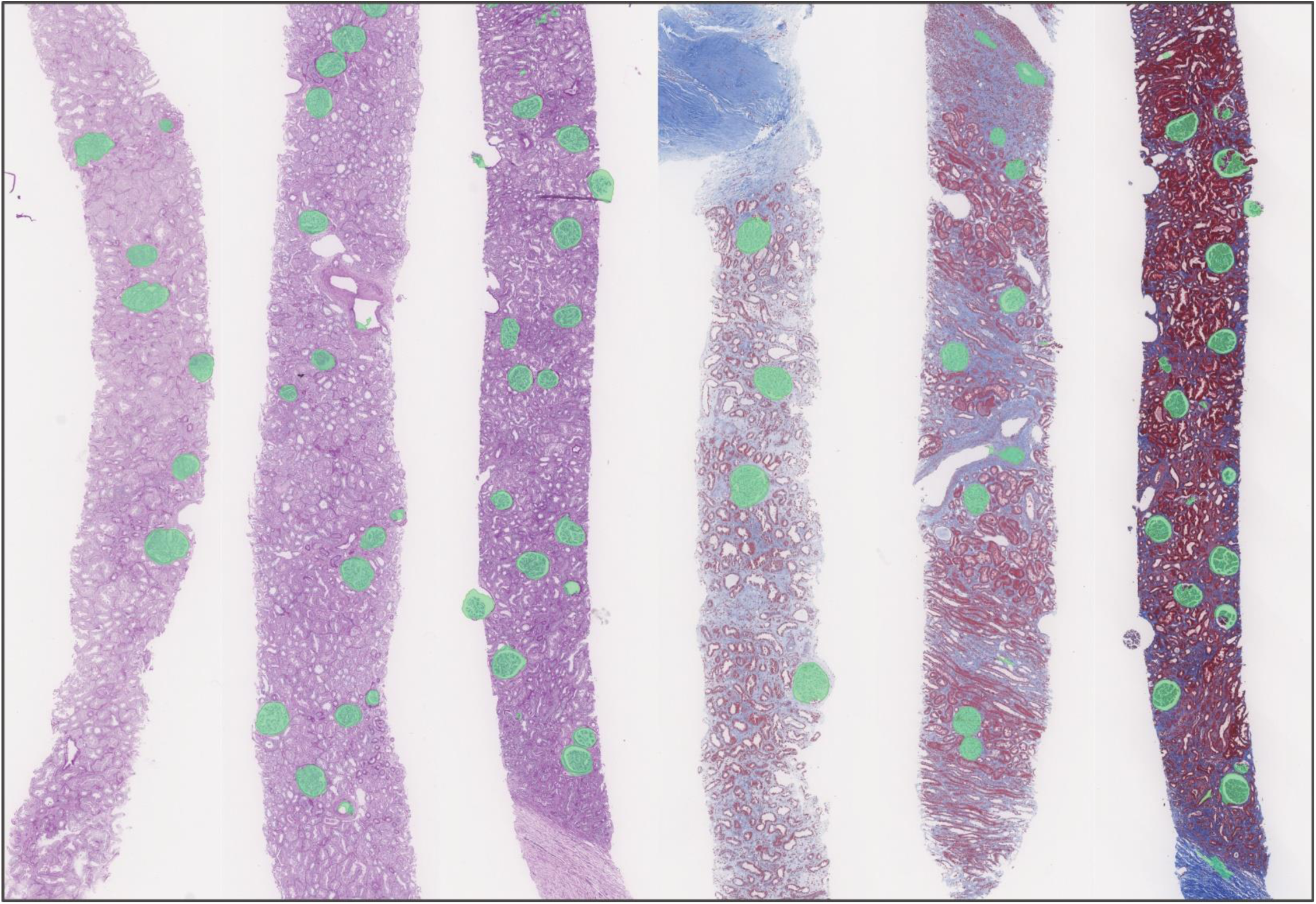
Examples of holdout segmentation using the datasetGAN training set. Examples of glomeruli segmentation shown in holdout WSIs. These segmentations are predictions by the DeepLab v3+ network, trained using a training dataset of 50,000 images that were generated using datasetGAN.

## DISCUSSION

The application of *datasetGAN* for the segmentation of histology tissue slides is very promising. The datasets used to test *datasetGAN* in the original paper [14] contained objects that were centered in the field of view such as faces or cars. Our training set of histology data used to train *datasetGAN* contained randomly cropped patches from WSIs, making structures such as glomeruli not centered and not always present in the training field of view. Despite this, *datasetGAN* seemed to do a very good job producing synthetically annotated histology datasets. Our tests show the performance of a model trained using *datasetGAN* synthetic images outperforms models trained with a limited number of traditional annotations. In our experiments we found that *datasetGAN* easily outperforms traditional training even when more than six times the amount of annotation effort is used to produce the training set.

Perhaps the most promising result was the ability of *datasetGAN* to improve a model when used for transfer learning [17]. Even if the desired segmentation performance is not achievable using *datasetGAN* annotations alone, our results show that the model created using *datasetGAN* and transfer learning has drastically improved performance when compared to hand annotation alone. This is particularly interesting when considered in the context of human in the loop annotation. In our previous work H-AI-L [6] we showed that human in the loop annotation greatly eases the amount of annotation effort required to develop a performant segmentation model. *DatasetGAN* makes a perfect addition to the H-AI-L method, and could be used to reduce the amount of initial annotation, accelerating the annotation process in future iterations.

While the *datasetGAN* reduces the amount of human annotation effort, it is worth noting that training the *styleGAN* model to produce realistic looking renal tissue patches took over 2 weeks of compute time using 2 Nvidia Tesla Quadro P100 GPUs. However, once the base *styleGAN* model was trained, further training of *datasetGAN* to produce glomeruli annotations only took a few hours using the 24 annotated synthetic patches. We have released this trained *styleGAN* model which can be used to train *datasetGAN* for any renal tissue segmentation task by the community. In the future we would like to explore using *datasetGAN* to accelerate the development of a model for nuclei segmentation from renal tissue biopsies. We also note that the training time for the *styleGAN* network could be reduced significantly with access to more computational hardware, specifically GPUs.

We believe that the performance of the segmentation model developed with *datasetGAN* annotations could have been improved with access to more capable computational hardware. To meet the memory requirements of the *datasetGAN* training code we were only able to produce synthetic annotations of size 256×256 pixels, four times smaller in each dimension than the patches of 1024×1024 that *styleGAN* is capable of producing. An example of this resolution mismatch can be observed in Figure 1, comparing the *styleGAN* outputs and *datasetGAN* outputs. Effectively this meant that the synthetic data produced by *datasetGAN* used to train the *deepLab v3+* network was a lower resolution, with a smaller field of view than the data used to train the *control* and *control+* models. Additionally, the both *control* models were trained using our *Histo-Cloud* software [4] which includes a highly optimized input pipeline for training d*eepLab* using WSI data, including image augmentation and extraction of image patches across multiple resolutions. We originally trained the *control* and *control+* models using 256×256 pixel patches, down sampled from the native slide resolution by four times (to match the *datasetGAN* synthetic dataset) and the performance of these models was lower than using larger patches. This performance gap suggests that higher resolution synthetic images would also benefit training for the *datasetGAN* model. In the future we would like to use a more capable system to optimize *datasetGAN* to produce training data at a higher resolution and see if this improves the performance of the downstream *deepLab* model.

## METHODS

### StyleGAN training

The first step to using *datasetGAN* to produce synthetic segmented images is training the base GAN model used to generate the images. For this *styleGAN* was trained using 2455 renal biopsy WSIs scanned at 40X magnification. This dataset came from a variety of institutions and included PAS, H&E, and Trichrome staining. To efficiently accomplish this training using such a large dataset, we used our Histo-fetch pipeline [18] to randomly select patches from the WSIs on the fly during training. *StyleGAN* was trained using 2 Nvidia Quadro P5000 RTX GPUs for 17,560,000 steps using a learning rate of 0.003 for the final resolution level 1024×1024. Further details of the *styleGAN* training are available in the ffhq standard configuration documented in the code. Our version of *styleGAN* that has been modified to train on WSIs is available here: https://github.com/SarderLab/WSI-stylegan.

### DatasetGAN training

To train *datasetGAN* to produce fake images, a limited number of synthetic images needed to be labeled by hand. To do this, the previously trained *styleGAN* was used to produce a large number of synthetic renal tissue patches. 24 of these patches containing glomeruli were selected and the glomeruli locations were annotated. Our *styleGAN* model trained on renal histology slides and the 24 annotated images used to train *datasetGAN* are available here: https://buffalo.box.com/s/t9qeitp3frxzmniaeq552b3b4cajbynf. To train *datasetGAN* using the hardware available to us, we reduced the resolution of the *styleGAN* output to 256×256 pixels. This allowed us to train the *datasetGAN* encoder to produce fake annotations corresponding to the generated images. The version of the *datasetGAN* code we used to perform this training is available here: https://github.com/SarderLab/datasetGAN_release. To generate the final training set used to train the *deepLab* segmentation network, we used our *datasetGAN* model to generate a training set of 50,000 image patches and annotations.

### DeepLab training

All the experiments conducted in this paper were trained using the *deepLab V3+* architecture [16]. To train the model using the 50,000 synthetic patches produced by *datasetGAN* we used the official implementation of *deepLab* and encoded the patches into tfrecord files for training. This training was done using 2 Nvidia Quadro P5000 RTX GPUs with a batch size of 12, and polynomial learning rate of starting at 0.007, for 30,000 steps. The training used a segmentation model pretrained on the ImageNet dataset [19] for transfer learning.

Training the *control* and *control+* models was done using *Histo-Cloud* [4] which extracts patches on the fly from annotated WSIs during training. Training was done using the same hyperparameters as the *datasetGAN* model training (described above), and also used the ImageNet model for transfer learning. The patches used for training were 512×512 and were extracted from the WSIs at a variety of downsampled scales including 1X, 2X, 3X, and 4X downsampled with respect to the native slide resolution. The details of the Histo-Cloud method of on the fly patch selection are further described in [4].

The *transfer learning* model was trained by repeating the training using the *control* dataset, but instead of initializing the network parameters using the ImageNet pretrained model, parameters from the *datasetGAN* trained model were used.

### Segmentation datasets

The dataset used generated by *datasetGAN* included 50,000 synthetic image patches that were 256×256 pixels at an effective 10X downsampled resolution with respect to the 40X WSIs used for the other training. The *control* dataset contained four WSIs containing 76 annotated glomeruli. This dataset contained 2 PAS and 2 Trichrome slides, and glomeruli were hand annotated. The *control+* dataset added 5 additional WSIs to the control set including 3 Trichrome and 2 H&E stained slides. All the slides in the control datasets were from a simple institution. The holdout dataset contained 23 slides, with 10 PAS, 8 Trichrome, and 5 H&E stained WSIs. The majority of these slides were selected from the same institution as the control training sets but one from each stain originated from an separate institution. In total the holdout dataset contained 486 hand annotated glomeruli.

## ACKNOWLEDGEMENTS

This project was supported by NIH-NIDDK grant R01 DK114485 (PS), NIH-OD grant R01 DK114485 03S1 (PS), a glue grant (PS) of the NIH-NIDDK Kidney Precision Medicine Project grant U2C DK114886 (Contact: Dr. Jonathan Himmelfarb), and NIH-OD grant U54 HL145608 (PS).

We would like to thank Kuang-Yu Jen for providing a large portion of the slides used for the control and holdout datasets.

## AUTHOR CONTRIBUTIONS

B.L. wrote the manuscript, the code, and performed the experiments. P.S. conceived the overall research plan and edited the manuscript.

## ADDITIONAL INFORMATION

The authors declare no competing interests.

